# RUPEE: A fast and accurate purely geometric protein structure search

**DOI:** 10.1101/475301

**Authors:** Ronald Ayoub, Yugyung Lee

## Abstract

Given the close relationship between protein structure and function, protein structure searches have long played an established role in bioinformatics. Despite their maturity, existing protein structure searches either use simplifying assumptions or compromise between fast response times and quality of results. These limitations can prevent the easy and efficient exploration of relationships between protein structures, which is the norm in other areas of inquiry. We have developed RUPEE, a fast, scalable, and purely geometric structure search combining techniques from information retrieval and big data with a novel approach to encoding sequences of torsion angles.

Comparing our results to the output of mTM, SSM, and the CATHEDRAL structural scan, it is clear that RUPEE has set a new bar for purely geometric big data approaches to protein structure searches. RUPEE in top-aligned mode produces equal or better results than the best available protein structure searches, and RUPEE in fast mode demonstrates the fastest response times coupled with high quality results.

The RUPEE protein structure search is available at http://www.ayoubresearch.com. Code and data are available at https://github.com/rayoub/rupee.

## Introduction

Proteins represent the functional end-product within the central dogma of molecular biology [1]. As such, understanding protein structure is a central goal within structural bioinformatics. Protein structure determination, prediction, alignment, and search all serve to advance this understanding. Below, we present our approach to a fast, scalable, and purely geometric protein structure search we refer to with the acronym of *RUn Position Encoded Encodings* of residue descriptors (RUPEE).

Given a protein domain identifier, whole chain identifier or an uploaded PDB file, RUPEE can search for matches among domains defined in SCOPe 2.07 [2], CATH v4.2 [3], ECOD develop210 [4], or among whole chains defined in the PDB. RUPEE is able to search either of these databases using any identifier. For instance, you can search SCOPe using a CATH domain identifier.

RUPEE has two modes of operation, fast and top-aligned. Fast mode is significantly faster than all other protein structure searches discussed below but at the expensive of accuracy. Despite this, we will show that the accuracy of RUPEE in fast mode is not far below that of the best available structure searches. On the other hand, the accuracy and response times of RUPEE in top-aligned mode are comparable to currently available protein structure searches that are commonly considered fast.

RUPEE stands out as not just another protein structure search, of which there are many. RUPEE is the first, to our knowledge, purely geometric protein structure search to achieve results as good as the best available protein structure searches. Additionally, it can be argued that the kinds of matches that RUPEE does return have more added value than the current state of the art in that with equal scores it is able to return results not biased toward a structure classification hierarchy such as SCOPe or sequence clusters such as the PDB-90. In this regard, RUPEE makes a fundamental contribution to protein structure research that lends itself to being leveraged in existing systems. It also provided a path for further research activity in the direction of big data representations of protein structures.

Besides our approach to protein structure search, we introduce a polar plot for torsion angles that may have wider applicability in the study of protein structure. Further, the *run position encoding* heuristic introduced below may have wider applicability to algorithms for character sequences containing long runs of repeats.

We first discuss some related work to provide a context for our approach followed by a description of our method. We end with a comparison of results against the mTM-align structure search [5], the secondary structure matching (SSM) search [6], and the CATHEDRAL structural scan [7] available at the CATH website.

## Related work

Pairwise alignment involves finding a set of spatial rotations and translations for two protein structures that minimizes a distance metric. Most commonly, the root mean squared deviation (RMSD) between *α*-carbons of aligned residues is minimized.

The typical use case of aligning one protein structure to another does not impose tight response time requirements. For this reason, pairwise alignments can focus on accuracy. On the other hand, a protein structure search can involve thousands of comparisons and accuracy is often balanced against speed. In this case, pairwise alignment is still useful for evaluating the results of a search, and this is the approach we take.

For pairwise alignment, Combinatorial Extensions (CE) [8] and FATCAT [9] are among the most popular tools, representing rigid and flexible protein alignments, respectively. CE performs a rigid alignment in order to minimize RMSD and FATCAT allows for a constrained number of twists in the protein chain in order to find a more flexible alignment before minimizing RMSD.

Whereas pairwise structure alignments only depend on the sequence of *α*-carbon coordinates, protein structure searches often introduce a further dependence on the sequence order of amino acids. This approach often takes the form of clustering proteins based on sequences and pre-calculating results for pairwise alignments among cluster representatives. Then, these pre-calculated results are used for filtering the number of structures used for comparisons against a query protein. The exact formula for combining the use of representatives and pre-calculated results varies from system to system. However, all systems using this approach share the same disadvantage, an indirect dependence on amino acid sequences. In the absence of a reliance on sequence representatives and pre-calculated results, and without sacrificing accuracy, response times suffer greatly, often taking upwards of an hour for queries to complete.

For protein structure searches, VAST [10] and the FATCAT server [11] are among the most popular. Nonetheless, these searches are slow in comparison to mTM, SSM, and CATHEDRAL when pre-calculated results are not used. If given a known protein domain, VAST can return structural neighbors in seconds using pre-calculated results. However, if uploading a PDB file where pre-calculated results are not used, response times for VAST can exceed 30 minutes. Similarly, the FATCAT server, that does not use pre-calculated results, can take over an hour to send results for a search against PDB-90 representatives [12].

Given the above, there remains a need for a purely geometric protein structure search. For the serendipitous exploration of relations between protein structures performed in the trenches, this search should be fast. Moreover, with a 10% yearly growth rate of solved structures deposited in the PDB [13], this search should be scalable. At a minimum, RUPEE takes a significant step in this direction as will be shown below.

## Methods

Broadly, we define a linear encoding of protein structure and convert this linear encoding into a bag of features. Min-hashing and locality sensitive hashing (LSH), techniques drawn from big data, are then applied to implement a protein structure indexing method that serves as the foundation for both RUPEE operating modes, fast and top-aligned.

Protein structure searches that use linear encodings are not unique [14–16]. The novelty of our approach lies in its remarkable performance given its simplicity. Additionally, elements of our approach can be isolated and found to be useful in their own right.

### Regions of Torsion Angles

Our first step towards a linear encoding of protein structure is to identify separable regions of permissible torsion angles, but first we introduce a new plot of torsion angles better suited to this effort.

Despite their utility and familiarity, Ramachandran plots [17] represent angular data using a square plot better suited for scalar data. This leads to the unwieldy arrangement where the top part of the plot is continuous with the bottom and the left is continuous with the right.

To identify regions of torsion angles, we randomly sampled 10,000 residues from high-resolution CATH s35 representatives to account for precision and redundancy, respectively. A Ramachandran plot of the sampled torsions angles is shown in the left plot of Fig 1. As can be seen, a single cluster of residues, consisting primarily of *β*-strands, appears at all 4 corners of the Ramachandran plot.

**Fig 1.**
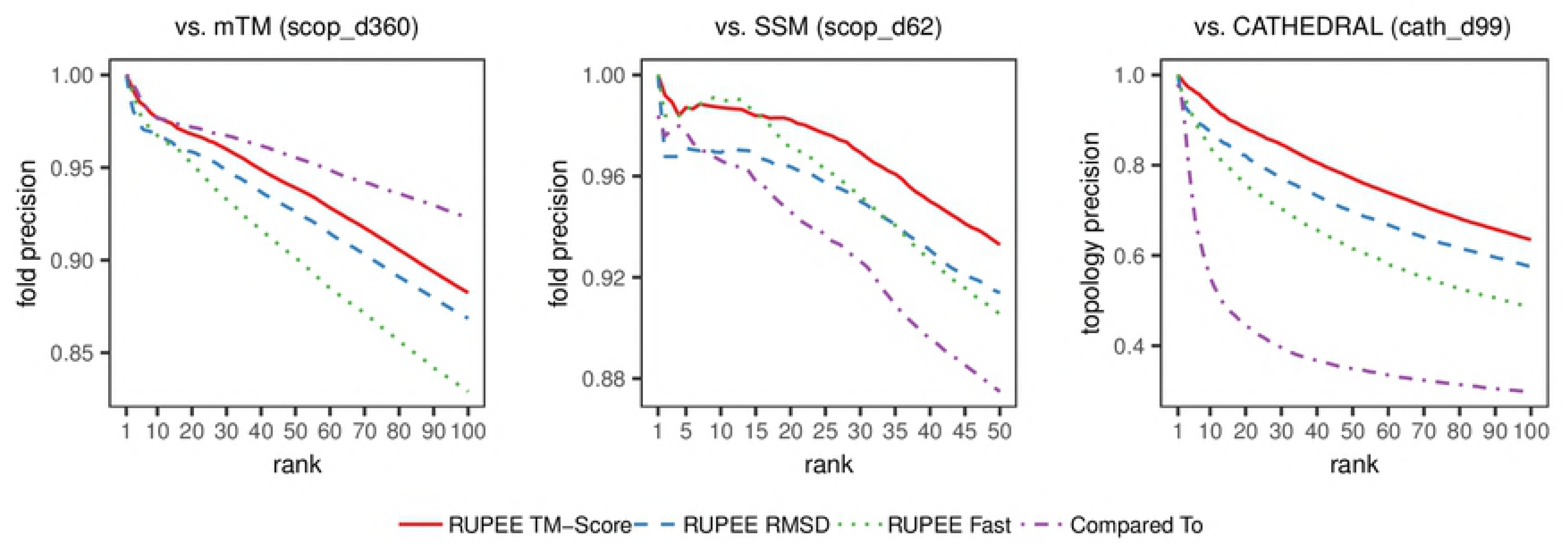
Ramachandran plot (right) and polar plot (left) of randomly sampled torsion angles

This continuity problem was partially addressed in [18] using *wrapped* and *mirrored* plots. Both wrapped and mirrored plots take advantage of the sparsely populated areas of the Ramachandran plot at *φ* = 0° and *ψ* = −120°. However, with larger samples of torsion angles, the area at *ψ* = −120° becomes less sparse. The use of a polar plot resolves this elegantly by only requiring one break in continuity at *φ* = 0°.

In the right plot of Fig 1, we show the same torsion angles appearing in the Ramachandran plot using a polar plot. In this plot, *φ* corresponds to the radius *r* and *ψ* corresponds to the angle *θ* in traditional polar plots. Notice the residues appearing at the 4 corners of the Ramachandran plot now appear in one continuous region of the polar plot centered at *φ* = *±*180° and *ψ* = *±*180°.

### Linear Encoding of Protein Structure

The polar plot described above is used to define torsion angle regions for each secondary structure assignment. The eight DSSP secondary structure assignment codes defined in [19] divide into three groups in which torsion angle regions are roughly the same: ‘G’,‘H’,‘I’, and ‘T’ corresponding to 3_10_-helix, *α*-helix, *π*-helix, and turn, respectively; ‘E’ and ‘B’ corresponding to *β*-strand and *β*-bridge, respectively; and ‘S’ and ‘C’ corresponding to bend and coil, respectively.

Polar plots for each group of DSSP assignment codes along with defined region and descriptor designations are shown in Fig 2, with the exception of turns and bridges, which receive descriptors 11 and 12, respectively. For each polar plot, there are well-defined continuous regions of torsion angles that remain continuous in the plots. The only exception is found in the bends and coil plot at *ψ* = 60° between *φ* = −180° and *φ* = 0°.

**Fig 2.**
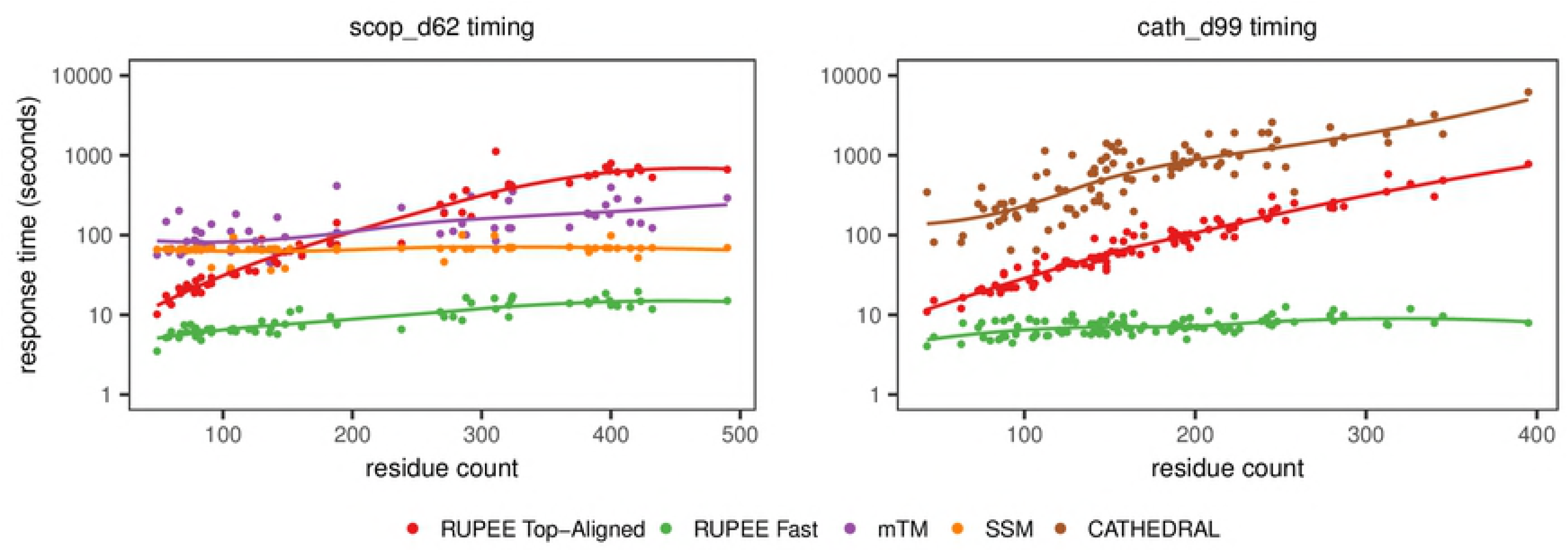
Polar plots of randomly sampled torsion angles with designated descriptors for region and DSSP code combinations

As an example of our linear encoding, we apply our method to the *β*-turn-*β* motif shown in Fig 3. The corresponding sequence of residue descriptors is shown below.

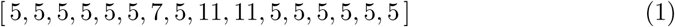

**Fig 3.**
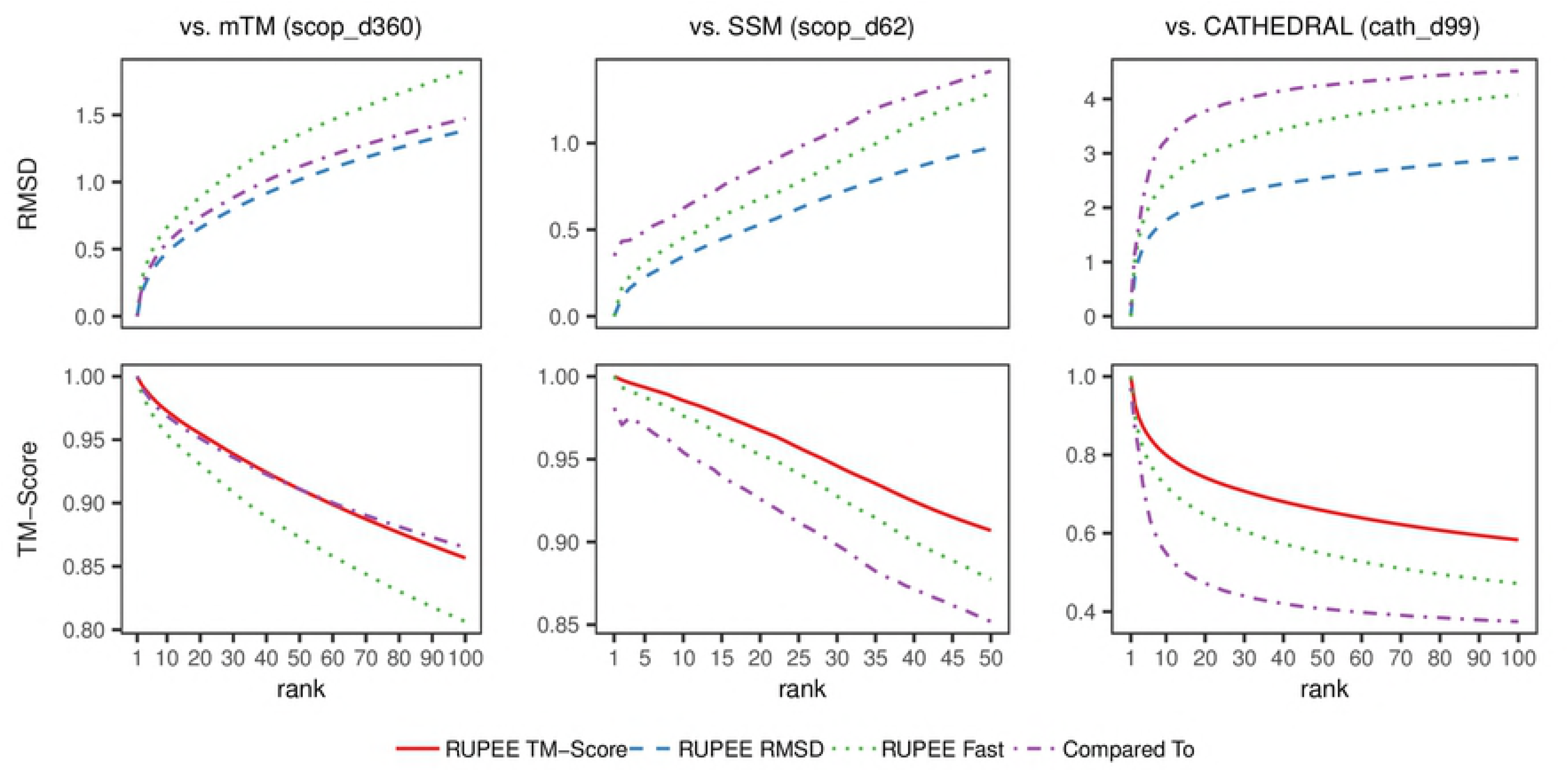
*β*-turn-*β* motif from CATH domain 1nycA00

### Bag representation of protein structure

Once a linear encoding for a protein structure is obtained, it needs to be further transformed into a representation suitable for fast and scalable similarity comparisons to other structures. The processing of text documents within Information Retrieval (IR) has long been used to satisfy these requirements using bag representations. There are two distinct categories of representations for documents, syntactic and semantic, and much of the research applying IR to protein structure search has focused on the latter [20–22].

We have adapted the syntactic approach to document similarity, often referred to as shingling [23], to our linear encoding of protein structure. We transform a linear sequence of descriptors into a multiset of shingles consisting of 3 consecutive descriptors. The overlap between shingles ensures some of the order information within the original sequence is preserved in the bag.

The length of a shingle is chosen to balance false positives, in the case of shorter shingles, against false negatives, in the case of longer shingles. In [24], we used 4 consecutive descriptors for our shingles. With the additions discussed in the *Operating modes* section below, we have found RUPEE is more tolerant of false positives and so accordingly we have cast a wider net by decreasing the shingle length to 3.

By shingling, we obtain a multiset of ordered lists from an ordered list of numbers. As an example, the sequence in (1) is transformed into the following bag of shingles.

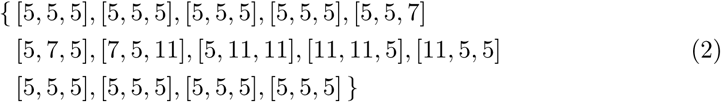

Next, each shingle *s* is hashed to an integer as shown in (3). The hash function used is a simplification of the hash function used in the Rabin-Karp algorithm [25]. The prime number 13 is used as the base since it is large enough to spread the descriptor values out in hash space without collisions.

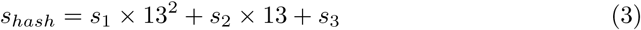

Subsequently, the multiset in (2) becomes the following bag of integers.

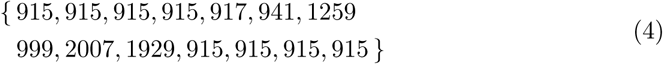

This final step completes the transformation of an ordered list of descriptors to a multiset of integers that still retains some of the order information present in the original list.

Notice in (4) the value 915, corresponding to the shingle [5, 5, 5], occurs frequently indicating the presence of *β*-strands. Since most proteins are dominated by regular secondary structure, the abundance of shingles for *β*-strands as well as the three types of helices, end up dominating comparisons. Moreover, since shingles are limited in length, this situation allows for structures with many short *β*-strands to match structures with fewer long *β*-strands. The same situation applies to helices.

To address this lack of specificity, we introduce a heuristic we call *run position encoding* (RPE). To distinguish between short and long runs, thereby increasing the specificity of the shingles, we add a factor of 10^5^ to each shingle hash as a function of the first residue’s position in a run *i*.

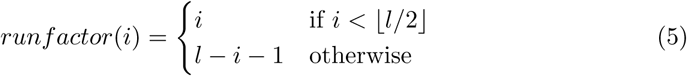

where *i* is zero-based and *l* is the length of the run. Multiplying by 10^5^ places the run factor as the left-most digit in the hash to avoid interference with the digits provided by the hash in (3). This placement is also convenient for visual inspection, since the run factor is isolated as the left-most digit.

The run factors for the sequence in (1) are

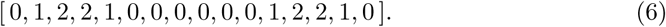

Applied to the bag of integers in (4) gives

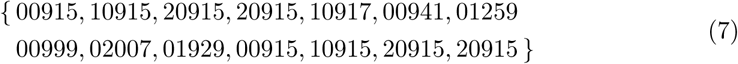

where the leading zero run factors are shown for clarity.

This pyramidal approach preserves matches at the boundaries between secondary structure runs and loops that would not otherwise be preserved in the presence of differences in run lengths of one or more.

Now that we have a representation of a protein structure as a bag of integers, similarity between any two structures *a* and *b* is defined as the Jaccard similarity [26] for multisets,

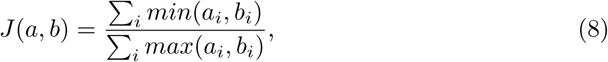

where *i* ranges over all possible shingle hashes *s*_*i*_ and *a*_*i*_ and *b*_*i*_ give the counts of shingle hash *s*_*i*_ in structures *a* and *b*, respectively.

### Min-hashing and LSH

In IR, the bag of shingles representation of documents is used in the near dupe clustering of documents [27]. One application of near dupe clustering is in the review stage of Electronic-Discovery [28], which is the most expensive stage in a discovery process. Often millions of documents must be examined by a staff of attorneys to make a reasonable effort at providing all documents relevant to the discovery request.

Grouping documents into near dupe clusters and assigning all documents within a cluster to a single reviewer reduces duplication of effort.

In the case of near dupe clustering, each document must be compared to every other document in the collection, taking quadratic time. For this task, min-hashing [29] and locality sensitive hashing (LSH) [30] can be combined to reduce this to subquadratic time. Although we do not near dupe cluster domains, we can still leverage the techniques of min-hashing and LSH to speed up protein structure search by a large constant factor.

Min-hashing is used to randomly select items from a set of items by repeatedly randomly hashing the items, sorting the hashes into a list, and then selecting the minimum item in each permuted list. If the same random permutation of items is performed on each set of items in a collection, the key result is that the probability of matching min-hashes is equal to the Jaccard similarity [29]. In order to approximate the Jaccard similarity for a given pair of sets, a sufficient number of min-hashes must be obtained.

In our case, the items are bags of shingle hashes for protein structures from which we obtain 99 min-hashes as described in [31]. Given the key result above, the Jaccard similarity can now be approximated by the proportion of matching min-hashes.

Next, we use the LSH banding technique as described in [31]. The key result of the banding technique is that if *any* band positions are a match for a given pair of structures, the probability that a specific similarity threshold has been met can be calculated. We use 33 bands of 3-min-hashes where the probability of a Jaccard similarity of 60% or greater is approximately 99%. Banding allows the problem of finding similar items to be parallelized across bands since all that is needed for a match is a single band match.

Together, min-hashing and LSH provide the foundation for both operating modes of RUPEE, fast and top-aligned.

### Operating modes

RUPEE provides two modes, fast and top-aligned. Each operating mode builds on the results provided by the min-hashing and LSH system described above. When a structure search is executed in either mode, a number of concurrent tasks are executed corresponding to the number of bands used for LSH. These concurrent tasks identify candidate matches based on a single band match and then validate the matches based on a comparison of min-hashes.

In fast mode, the top 8,000 matches are obtained along with the original gram sequences defined for the structures. A further validation is done by performing a longest common subsequence (LCS) analysis of the matched gram sequences and adjusting the Jaccard similarity scores accordingly. This step accounts for possible gram matches among pairs of structures that are out of order since the min-hashing and LSH techniques themselves do not consider the order of the gram matches. The final step of fast mode is to sort the matches based on the adjusted scores and return the results.

Top-aligned is an additional step following fast mode. First, unoptimized CE alignments are performed on the top 2,000 matches obtained from fast mode. Then optimized CE alignments are performed on the top 400 matches and these are finally returned sorted either by RMSD or TM-Score. Top-aligned is a simple layer following fast mode that establishes RUPEE fast mode as an effective filtering method that contains in its top 2,000 results enough good matches to compete with the best available structure searches.

## Results

Protein structure searches can be evaluated using pairwise alignment scores or by comparison of results against the hierarchy of a protein structure classification database. The RMSD of aligned residues is widely used in evaluations but is not perfectly suited to full-length comparisons between structures since distances between unaligned residues are not factored into the score. On the other hand, the TM-score [32] takes all residues into account. Among protein structure classification databases for which corresponding structure searches exist, SCOPe [2] and CATH [3] are the most popular.

For our results, we have created 3 benchmarks, scop d360, scop d62, and cath d99, for pairwise evaluations to mTM, SSM, and CATHEDRAL, respectively. scop d360 is derived from the d500 benchmark used in [5] filtered for domains in SCOPe 2.07 for which mTM returns 100 or more results. Similarly, scop d62 is derived from the d500 benchmark filtered for domains defined in SCOP 1.73 for which SSM returns 50 or more results. In keeping with our description of RUPEE in [24], the cath d99 benchmark contains 99 superfamily representatives from the top 100 most diverse superfamilies defined in CATH v4.2 for which CATHEDRAL returns results in less than 12 hours.

We perform pairwise evaluations to ensure the fairness of our comparisons. First, for domain searches, SSM is working with the SCOP 1.73 database, so accordingly we operate RUPEE on SCOP 1.73 domains to ensure RUPEE does not have more domains to work with for scoring and precision evaluations. Second, mTM is updated to work with SCOPe 2.07 domain definitions but still retains domains from 2.06 that have since been redefined either through mergers or splits in 2.07. On the other hand, CATHEDRAL presents no such challenges but still requires a separate benchmark since it is working with a distinct hierarchy, CATH v4.2.

All benchmark definitions can be found in S1 Benchmarks.

### Scoring

Fig 4 shows average cumulative values for each ranked result averaged over all searches. Both RMSD and TM-score values are shown, provided as outputs from optimized CE pairwise alignments. A TM-score above 0.5 is a good predictor for whether or not two domains are in the same fold [33]. TM-scores greater than 0.17 are considered potentially meaningful whereas TM-scores less than 0.17 are considered to be due to random alignment [32].

**Fig 4.**
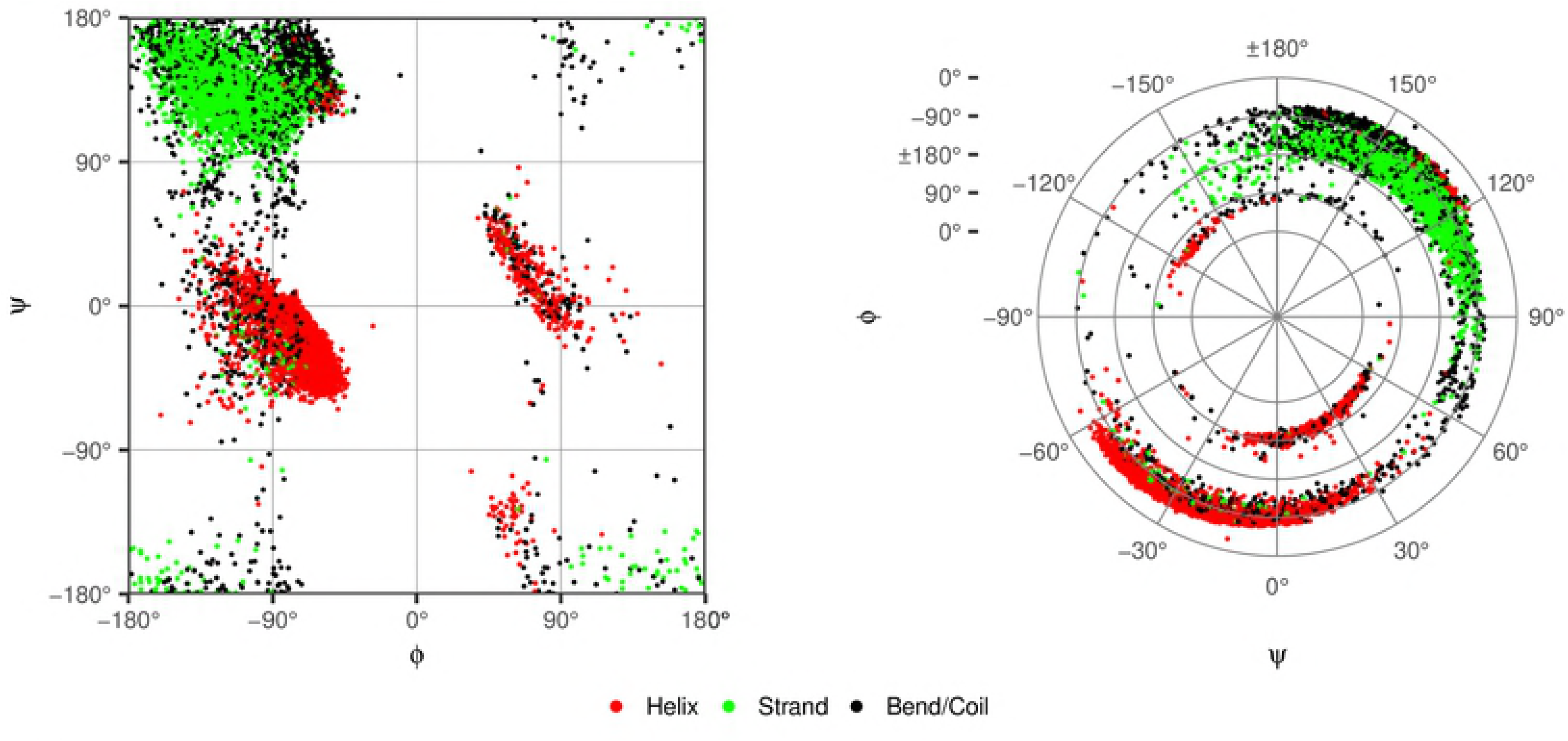
Scoring from CE pairwise alignments for RUPEE fast, RUPEE top-aligned sorted by TM-Score, and RUPEE top-aligned sorted by RMSD

A figure similar to Fig 4 but using FATCAT for pairwise comparisons can be found in S2 Fig. The same relative relationships hold with only small variations.

RUPEE fast, top-aligned sorted by RMSD, and top-aligned sorted by TM-Score, perform better than SSM and CATHEDRAL. The scoring in the cath d99 benchmark comparisons are notably lower than for the other two benchmarks. This is expected since CATHEDRAL only returns CATH s35 representatives. Likewise, for this comparison RUPEE is filtered for s35 representatives to match. Given that the cath d99 benchmark is evaluated against representatives, there are fewer highly similar structures returned in the results.

In our evaluation, mTM faired better than SSM and CATHEDRAL. mTM also performed better than RUPEE fast, although RUPEE fast is still within 0.08 TM-Score points of mTM at the 100^th^ result, which is notable considering its speed.

For TM-Score, RUPEE top-aligned and mTM are nearly identical with RUPEE slightly better for ranks less than 50 and mTM better for ranks greater than 50. For RMSD, RUPEE top-aligned does perform better than mTM but this can most likely be attributed to the fact that mTM only sorts by TM-Score. If mTM sorted by RMSD we suspect the results again would be nearly identical. Nevertheless, it is worth noting that the initial LSH and min-hashing technique does not explicitly bias results towards one particular measure.

### Precision

Fig 5 shows precision (i.e. positive predictive value or PPV) averaged over all searches, where positive results are defined as domains with the same classification for the indicated hierarchy level as the query domain. A plot of recall is unnecessary since Fig 5 provides precision at specific ranks for identical sets of searches. Hence, recall curves have the same relative relationships as those shown for precision.

**Fig 5.**
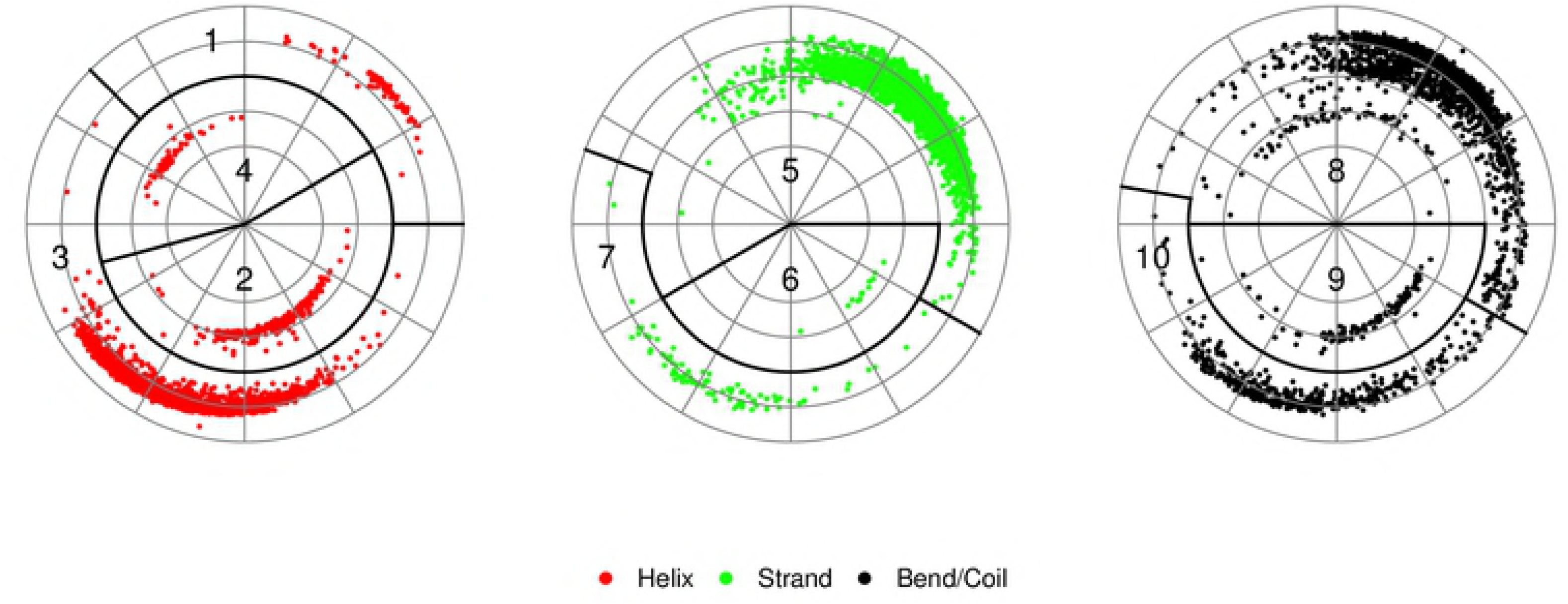
Precision for RUPEE fast, RUPEE top-aligned sorted by TM-Score, and RUPEE top-aligned sorted by RMSD

We should expect a structure search to have reasonable precision with respect to the hierarchy levels of the structure classification it is searching. However, it is not clear how to define reasonable. On the other hand, if precision is too high, the search provides little value beyond that provided by the structure classification hierarchy it is searching. Towards the extreme end of high precision, it would be sufficient for a search to return the best match and from there refer to the hierarchy for additional results.

Similar to the scoring evaluations above, RUPEE fast and top-aligned, sorted by both RMSD and TM-Score, show higher precision than SSM and CATHEDRAL with the exception of RUPEE top-aligned sorted by RMSD for ranks lower than 10 compared against SSM.

Again, mTM faired better than SSM and CATHEDRAL. Notably, mTM also shows clearly higher precision than RUPEE top-aligned, sorted by both RMSD and TM-Score. In the absence of Fig 4, one may be led to regard this as a negative result. However, to the contrary, in the presence of Fig 4, where it is shown that RUPEE and mTM have almost identical scoring, this result is remarkable. This disparity suggest that RUPEE is better able to find significantly similar matches not necessarily aligned with the SCOPe hierarchy. In fact, since the initial min-hashing and LSH technique used by RUPEE check for similarity to all available structures, no structure is able to hide behind a sequence based cluster representative. So this result is in keeping with how RUPEE is intended to work and how it is described above.

### Response Times

Fig 6 shows response times in seconds for the scop d62 and cath d99 benchmarks. Here, we are able to show RUPEE fast and top-aligned, mTM and SSM on the same plot because scop d62 is a subset of the scop d360 benchmark. Both plots are shown with a logarithmic scale in order to include all outliers while still being able to view the overall trends in response times. Loess regression curves are also provided to further highlight the overall trends.

**Fig 6.**
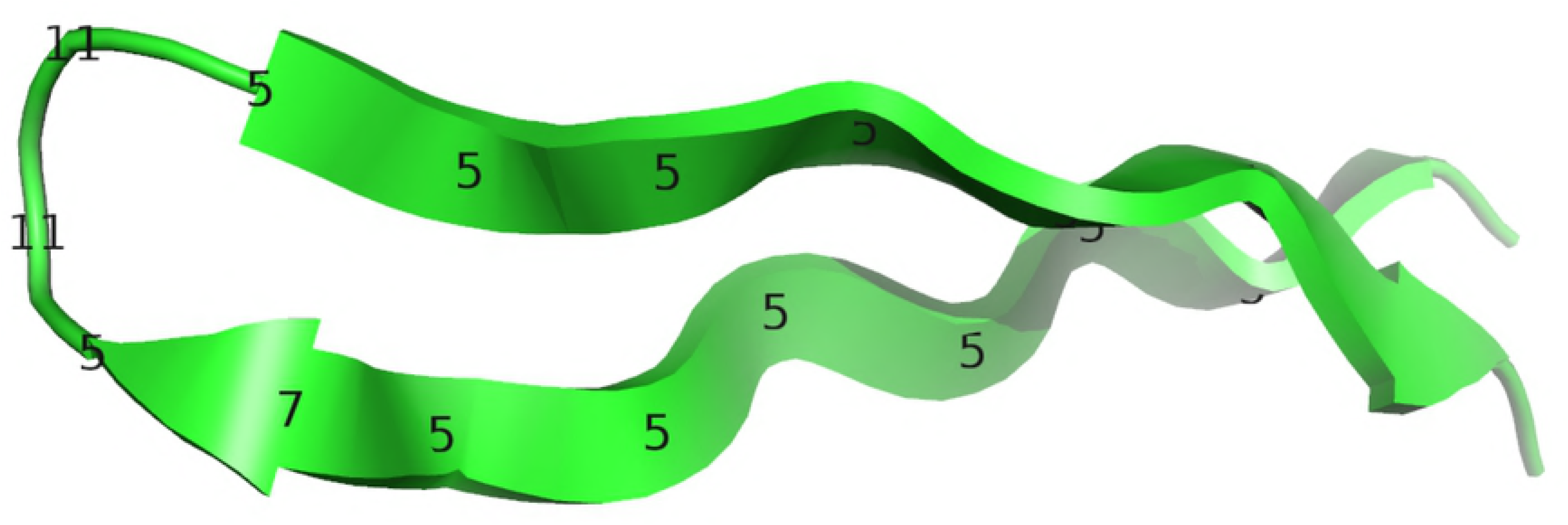
Response times for RUPEE fast and RUPEE top-aligned. The response times for RUPEE top-aligned are dominated by pairwise structure alignments and do not depend on the sort order.

In all cases, RUPEE fast is considerably faster than all other searches. It is also clear that fast mode is not as sensitive to increasing residue counts in contrast to CATHEDRAL and RUPEE top-aligned. Response times for SSM are not affected by residue counts at all and always returns results in less than 100 seconds but this is at the expense of performance as shown in Fig 4.

In the left plot of Fig 6, it is shown that RUPEE top-aligned is faster than mTM for residue counts below 200, but then response times for RUPEE increase beyond that of mTM. Down the stretch, mTM provides increasingly better response times than RUPEE, while RUPEE is still able to provide reasonable response times. The right plot of Fig 6 shows RUPEE top-aligned is significantly faster than CATHEDRAL for all residue counts.

The trend of increasing response times for RUPEE top-aligned is a direct result of the pairwise structure comparisons that are performed on the top 2,000 results provided by RUPEE fast mode.

Response times to a large degree are a measure of the amount of resources available to an application. In the case of RUPEE, our response times were gathered from RUPEE running on a fairly old laptop with a 2nd generation Intel^®^ Core™ i7-2720QM Quad-Core CPU and 8 GB of memory. With more resources the response times of RUPEE can be further improved since pairwise alignments can be run in parallel. For mTM, SSM, and CATHEDRAL, we gathered response times by automating their respective web sites using the Selenium WebDriver API.

## Discussion

As shown above, a purely geometric big data approach to protein structure search can compete with the best available protein structure searches. Nonetheless, there remain avenues for further improvement and investigation.

While our min-hashing and LSH scheme does allow for some flexibility in the size of matched protein structures, it is not specifically designed for containment searches. However, the initial min-hashing and LSH search does operate fast enough, within seconds, that it does present the possibility of executing multiple searches within the context of a single query. With more resources, we can distribute min-hash and band data across multiple compute units, with each data set representing a subset of chopped structure representations. With a 75% overlap of these subsets, RUPEE searches effectively become containment searches.

We have tried the overlapping subset idea and have found it to be effective. However, on a single compute unit, the time required is more than we are willing to accept for this first iteration of RUPEE.

One possible drawback of RUPEE is that in both fast and top-aligned modes, it only returns the top 400 results. Again, with more resources, this number can be increased, but there still remains a need for some kind of cut-off. Nonetheless, in most search tools, having to looking past the first few hundred results usually indicates an ineffective search strategy. To this end, RUPEE provides filters that can be used for traversing structure space more efficiently. For SCOPe, CATH, and ECOD you can instruct the search to only return domains that differ from the query structure at a chosen hierarchy level classification. Additionally, for CATH you can filter results based on hierarchy level representatives. Since RUPEE does not rely on sequence clusters, these kinds of filters are easy to implement, do not reduce the number of returned results, and allow for the discovery of unexpected structural similarities across classification hierarchies.

One area that stands out for possible improvement is our longest common subsequence (LCS) scoring adjustment to the results initially returned by RUPEE. While the LCS step has been shown to be effective, it is notable for its simplicity. A more complex step of validating the sequence of gram matches can take a form similar to the path extension algorithm used by CE. In this case, sequences of matches would only be extended when the difference of interresidue distances between gram pairs already in the sequence and a candidate pair to be added to the sequence fall below some threshold.

On the other hand, it would be interesting to see where further analysis of the initial results returned by RUPEE before LCS and any sort of order enforcement beyond that of the grams themselves could lead. For instance, topological permutations such as circular permutations, segment-swapping and changing secondary structures within homologous proteins are not uncommon [34]. The initial RUPEE min-hashing and LSH algorithm provides candidate matches along with matched grams. With some thought, an algorithm similar to FATCAT can be developed allowing for permutations in addition to twists.

As can be observed from considering Fig 4 and Fig 5 together, there are good domain matches aligned with the SCOPe hierarchy that mTM is able to find that RUPEE does not. Conversely, there are good domain matches not aligned with the SCOPe hierarchy that RUPEE finds and mTM does not. A comprehensive listing of these difference may find interesting similarities not previously known.

## Conclusion

With the growth rate of solved structures deposited in the PDB, the need for a fast and scalable structure search is growing. Using run position encoded shingles of residue descriptors combined with min-hashing and LSH, we have shown that RUPEE fast is able to provide good results in seconds running on an Quad-Core laptop. Currently, RUPEE fast is the fastest available protein structure search providing the demonstrated level of accuracy. For RUPEE top-aligned, we have shown that a purely geometric big data approach to protein structure search is able to produce results equal to or better than the current state of the art protein structure searches that variously depend on clustered sequences or protein structure classification hierarchies. Moreover, we have shown evidence that suggests the results from RUPEE top-aligned provide more added value by discovering high scoring protein structure matches not necessarily aligned with a particular protein structure classification hierarchy or hiding behind cluster representatives. The ability for RUPEE to quickly examine all structures among hundreds of thousands sets it apart as a tool that can be used for discovering previously undetected structural similarities.

## Supporting information

**S1 Benchmarks. Domains included in benchmarks used for evaluation.**

**S2 Fig. Scoring from FATCAT pairwise alignments**.

**S3 Methods Addendum. Run factors for shingles instead of descriptors**.

